# Stereoselectivity of Aminoacyl-RNA Loop-closing Ligation

**DOI:** 10.1101/2024.11.26.625528

**Authors:** Shannon Kim, Marco Todisco, Aleksandar Radakovic, Jack W. Szostak

## Abstract

The origin of amino acid homochirality remains an unresolved question in the origin of life. The requirement of enantiopure nucleotides for nonenzymatic RNA copying strongly suggests that homochirality of nucleotides and RNA arose early. However, this leaves open the question of whether and how homochiral RNA subsequently imposed biological homochirality on other metabolites including amino acids. Previous studies have reported moderate stereoselectivity for various aminoacyl-RNA transfer reactions. Here we examine aminoacyl-RNA loop-closing ligation, a reaction that ‘captures’ aminoacylated RNA in a stable phosphoramidate product, such that the amino acid bridges two nucleotides in the RNA backbone. We find that the rate of this reaction is much higher for RNA aminoacylated with L-amino acids than D-amino acids. We present an RNA sequence that near-exclusively captures L-amino acids in loop-closing ligation. Finally, we demonstrate that ligation of aminoacyl-L-RNA results in inverse stereoselectivity for D-amino acids. The observed stereochemical link between D-RNA and L-amino acids in the synthesis of RNA stem-loops containing bridging amino acids constitutes a stereoselective structure building process. We suggest that this process led to a selection for the evolution of aminoacyl-RNA synthetase ribozymes that were selective for L-amino acids, thereby setting the stage for the subsequent evolution of homochiral peptide and ultimately protein synthesis.

## Introduction

In all known life, nucleic acids and proteins are composed exclusively of D-nucleotides and L-amino acids, respectively. The universality of biological homochirality implies that the use of enantiopure nucleotides and amino acids was established very early in the origin of life.

Experiments with model systems for nucleic acid copying and catalysis show that heterochiral components impede reactions that are fundamental for the origin of life.^1,2^ Nonenzymatic RNA template copying reactions are severely inhibited when both D- and L-nucleotides are present.^3^ Furthermore, if RNAs containing both L- and D-nucleotides were produced, the copying of such heterochiral templates would likely fail due to inconsistencies in the orientations of bound substrates.^3^ Ribozyme catalysis, in the form of a polymerase ribozyme that can synthesize homochiral RNA from mixtures of L- and D-nucleotides,^4^ may appear to offer a solution, but the ribozyme itself must start out as homochiral RNA, suggesting that homochiral components had to precede the emergence of ribozymes. Homochiral nucleotides therefore appear to be a prerequisite for the emergence of genetic self-replication. However, it remains an open question whether homochirality in other classes of biomolecules, such as amino acids and lipids, was established as a consequence of the prior homochirality of RNA. For example, if early metabolic reactions were catalyzed by a network of ribozymes, the intrinsic stereoselectivity of macromolecular catalysis could have led to the stereoselective synthesis of chiral metabolic intermediates as well as products such as amino acids. Alternatively, the chirality of other biomolecules could, in principle, have been established independently.

Considerable research has gone into mechanisms of symmetry breaking with regard to the origin of life.^5,6,7,8,9^ The autocatalytic Soai reaction can yield near-homochiral alkylated pyrimidine 5-carbaldehydes from very slight enantiomeric imbalances in the starting material.^10^ Viedma ripening, through attrition-enhanced deracemization, can result in a homochiral solid phase of conglomerate crystals from a racemic solution.^11^ However, neither of these processes has been shown to result in the homochiral synthesis of prebiotically relevant compounds. A recent proposal for the origin of biological homochirality leverages the chiral-induced spin selectivity (CISS) effect to achieve stereoselective crystallization on a magnetically polarized surface. Tassinari et al. first reported the separation of the enantiomers of the amino acids asparagine and glutamic acid by stereoselective crystallization.^12^ However, crystallization-based approaches to the chiral resolution of amino acids face the issue that only four biological amino acids (asparagine, aspartic acid, glutamic acid, threonine) have been observed to crystallize as conglomerates.^12,13^ Furthermore, conglomerate crystals of aspartic acid and glutamic acid are formed only when crystallization proceeds from a supersaturated solution.^14^ The use of magnetic polarization for enantioselective crystallization has found greater success as a potential origin of homochirality in nucleotides. In a study by Ozturk et al., enantiospecific nucleation of crystallization of riboseaminooxazoline (RAO) was achieved via crystallization on a polarized magnetite surface, with L-/D-selectivity dependent on the direction of magnetic polarization.^15^ RAO is a nucleotide precursor in the cyanosulfidic prebiotic chemical network proposed by Xu and Sutherland.^16^ A process leading to homochiral RAO is particularly satisfying as the homochirality of RAO would lead directly to the homochiral synthesis of all downstream nucleotides, and thus to homochiral RNA and DNA.

Based on the biological pairing of D-RNA and L-amino acids, it is possible that the initial selection of L-stereochemistry for amino acids may have been governed by interactions with homochiral RNA. Previously, Tamura and Schimmel reported four-fold chiral selectivity for L-amino acids in aminoacyl transfer from an RNA 5′-phosphate to the 3′-hydroxyl of an upstream oligomer in a nicked duplex.^17^ Wu et al. showed up to 10-fold preference for L-amino acids in interstrand aminoacyl transfer from the 5′-phosphate to the diol of RNA.^18^ Roberts et al. demonstrated 5-to 10-fold higher yields at room temperature (and 10 to 50-fold at -16 ºC) for the formation of L-vs D-aminoacyl-ester RNA in a reverse loop-closing ligation beginning with the amino acid anchored to the 5′-phosphate as a phosphoramidate.^19^ On the other hand, Kenchel et al. examined the stereoselectivity of ribozyme-catalyzed self-aminoacylation and found that different ribozymes exhibited stereoselectivity for either D- or L-amino acids.^20^ We have recently described a two-step reaction system for the formation and capture of aminoacylated RNA in which the addition of activated amino acids to RNA results in aminoacylation followed by loop-closing ligation.^21,22^ Our results suggest that this process of aminoacyl capture could be stereoselective, but in that study we could not distinguish between selectivity at the stage of aminoacylation, ligation, or hydrolysis.

Here we report on the stereoselectivity loop-closing ligation with RNA that is pre-aminoacylated with either an L- or a D-amino acid. In this reaction the amine of an aminoacylated RNA attacks an activated 5′-phosphate to form a phosphoramidate linkage. We find a widely varying stereoselectivity in favor of RNA aminoacylated with L-amino acids for this reaction, with the magnitude of the selectivity depending on the RNA sequence and structure. With all RNA architectures, we observe a preference for the ligation of L-aminoacylated RNA, but with one structure in particular, we observe an almost 200-fold faster rate of loop-closing ligation when the RNA is aminoacylated with an L-amino acid, highlighting the impact of RNA tertiary structure on the stereoselectivity of this reaction. When we carry out aminoacyl loop closing ligation using L-RNA, we observe a reversed stereoselectivity in favor of D-amino acids, demonstrating that the handedness of RNA is the chirality determining factor in this system. Our results strengthen the hypothesis that stereoselection of L-amino acids occurred through chiral transfer from RNA, and more specifically, that the biological pairing of L-amino acids with D-RNA resulted from the stereochemical properties of aminoacylated RNA.

**Scheme 1..**
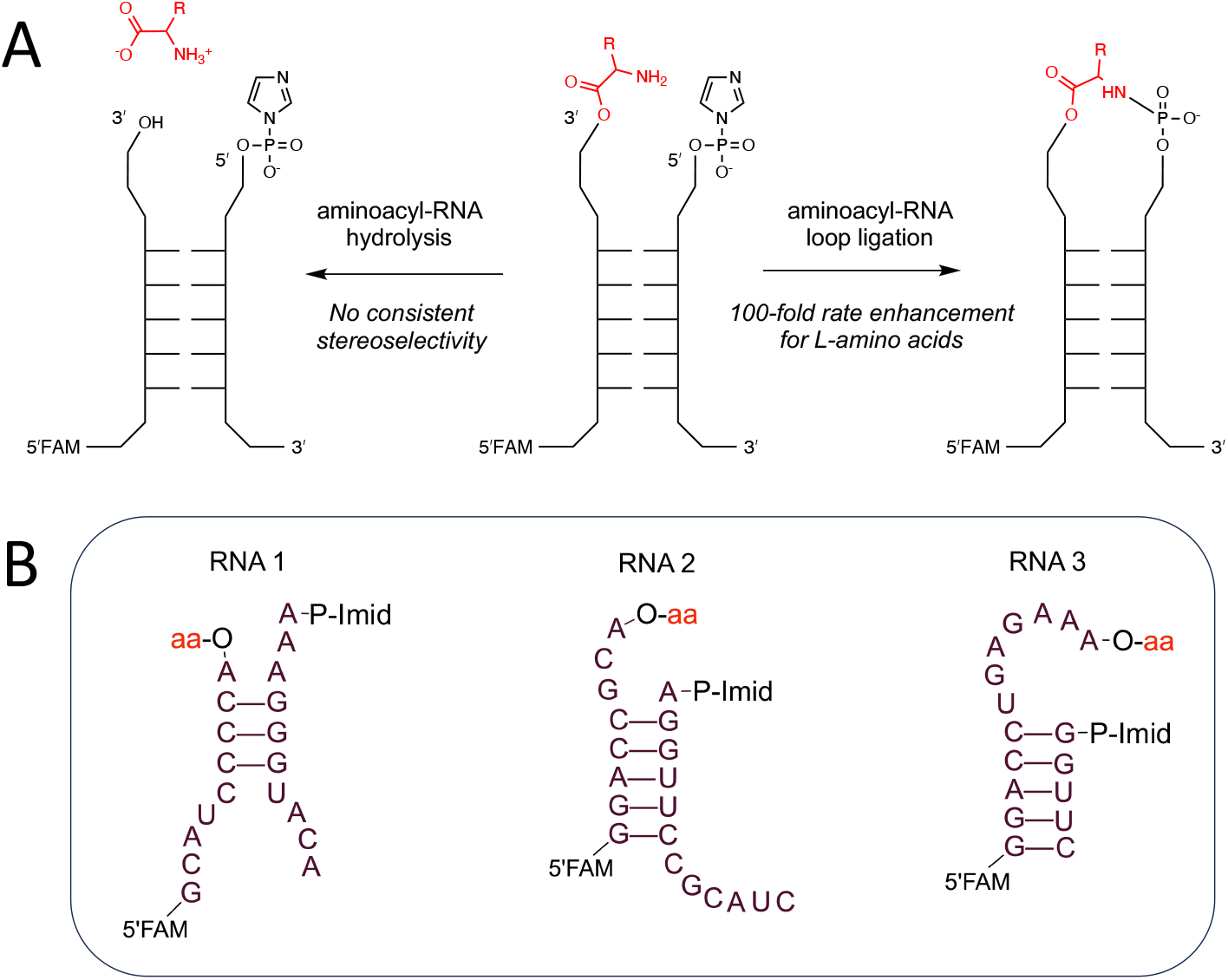
Outline of the reactions and RNA architectures studied in this work. **A**. Diagram of the reactions of aminoacylated RNA annealed to a capture strand with an activated 5′-phosphate (center). Left: hydrolysis of the aminoacyl ester. Right: ligation of the aminoacylated strand to the capture strand. Relative rates for L-over D-aminoacylated RNA reach the order of 10^2^, while hydrolysis of the aminoacyl ester occurs without a consistent stereochemical bias. **B**. Secondary structures of the three RNA architectures used in this study. P-Imid indicates activation of the nucleotide 5′-phosphate with imidazole.

## Results

Our previous studies of the aminoacyl capture reaction hinted at stereoselectivity at some stage of the process, which prompted us to investigate the stereoselectivity of this reaction in greater detail. For our experiments, we pre-aminoacylated the 2′(3′)-diol of an ‘acceptor’ RNA oligonucleotide using the Flexizyme, an aminoacyl-RNA synthetase ribozyme,^23^ which allows for RNA aminoacylation with both L- and D-amino acids.^24^ We then incubated the aminoacylated acceptor strand with a ‘capture’ strand carrying a 5′-phosphorimidazolide moiety. The acceptor and capture oligonucleotides anneal to form a duplex stem with non-complementary overhangs (**Scheme 1**), such that a successful ligation reaction produces an amino acid-bridged RNA stem-loop. We studied three different RNA architectures and measured the stereoselectivity of loop-closing ligation in each construct. Of the constructs, RNA 1 and RNA 2 (**Scheme 1**) were adapted from stem-loop domains of the Flexizyme ribozyme.

Architecture 3 (**Scheme 1**) emerged from a screen for self-aminoacylation with glycine followed by loop-closing ligation.^22^ As a preliminary test we examined the aminoacyl-RNA ligation of architecture 1 with L- and D-alanine. We observed greater loop-closing ligation with L-alanine, with 35 % of the L-alanylated acceptor strand ligating after 1 hour, as opposed to <1 % ligation of D-alanylated acceptor (**Figure 1A**).

**Figure 1.**
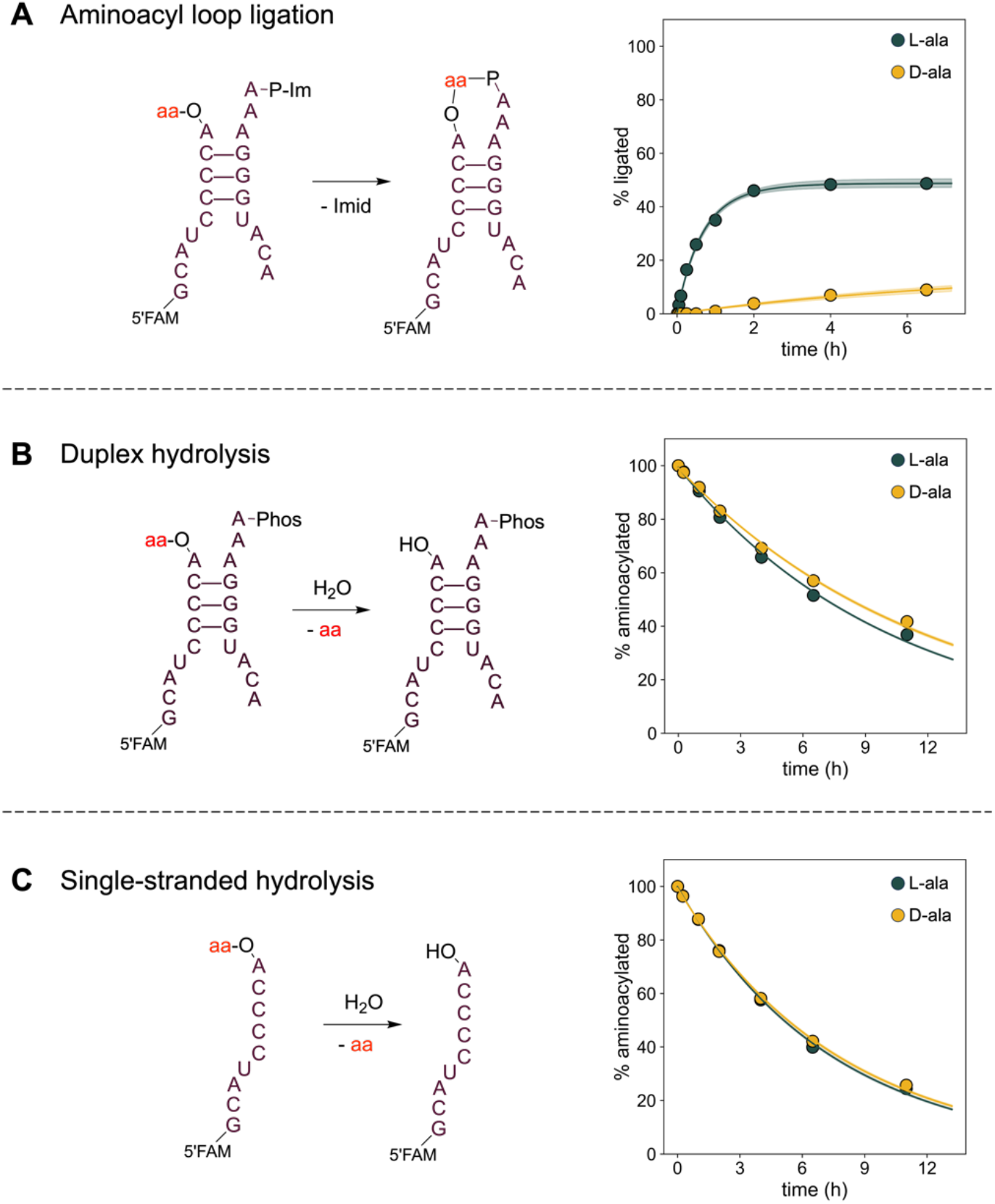
Reaction schemes and time courses for aminoacyl-RNA loop-closing ligation and hydrolysis. **A**. Left: reaction scheme for aminoacyl-RNA loop-closing ligation using an imidazole-activated capture strand. Right: time course of the ligation reaction. **B**. and **C**. Left: reaction schemes for aminoacyl-RNA hydrolysis under duplex and single-stranded conditions. Right: time courses of the hydrolysis reaction. Reaction rates are presented in Tables 1, S1 and S2, and gel images are shown in Figure S1. All reactions were conducted at 0 °C in 5 mM MgCl2, 100 μM Na2EDTA, 100 mM imidazole, pH 8.0 and 5 μM RNA oligonucleotides. Shaded regions envelope the 95% prediction interval as determined from the kinetic model.

To test whether the observed stereoselectivity might simply reflect stereoselective aminoacyl-RNA hydrolysis, we measured the rate of hydrolysis of the RNA-aminoacyl ester when the aminoacylated acceptor strand was annealed with a capture strand with an unactivated 5′-phosphate, so that ligation could not occur (we refer to this reaction condition as ‘duplex hydrolysis’). To account for any unanticipated effects of the capture strand on the stability of the aminoacyl-RNA, we also measured the hydrolysis rate in the absence of the capture strand (which we refer to as ‘single-stranded hydrolysis’). In the duplex condition, we observed that L-alanylated RNA hydrolyzed approximately 1.2 times faster than D-alanylated RNA (p<0.01, **Figure 1B**). In the single-stranded condition, the two stereoisomers hydrolyzed at approximately the same rate (**Figure 1C**), suggesting that the complementary strand may have a modest influence on the hydrolytic lability of the aminoacyl ester. When we performed these reactions with leucine, lysine and proline, we observed that that the two stereoisomers hydrolyzed at comparable rates (within a factor of 0.7 to 1.2-fold, **Tables S1-S2**), thus excluding preferential hydrolysis as the explanation for the stereoselective formation of the loop-closed product.

To explore the generality of the stereoselectivity of aminoacyl-RNA loop-closing ligation, we performed the ligation and hydrolysis reactions with three additional amino acids: proline, lysine, and leucine, chosen to represent a diversity of side chains. For all four amino acids, loop-closing ligation plateaued more quickly and led to higher yields with the L-enantiomer than with the D-enantiomer (**Figure 2**). In contrast there were minimal differences in the rate of hydrolysis between the two stereoisomers of the aminoacylated RNAs (**Tables S1–S2**). To quantitatively compare the loop-closing ligation kinetics with the two enantiomers of the amino acids, we kinetically modeled the ligation reactions with a set of differential equations to account for the aminoacyl-RNA hydrolysis (as determined from our experiments measuring hydrolysis in a duplex), and 5′-phosphorimidazolide hydrolysis (as determined from our recent work^25^). We also accounted for the reshuffling of oligonucleotides in complexes,^26^ since loop-closing ligation can only occur when both aminoacyl- and imidazole-groups are present on the oligonucleotide termini. In some cases, the maximum ligation yield was lower than expected based on the rate of aminoacyl-RNA hydrolysis (notably with Architecture 3, alanine and leucine). To account for this, we allowed the model to determine the initial percentage of imidazole-activation as a free parameter shared between the L- and D-aminoacylated constructs, since all pairwise experiments were conducted simultaneously. Upon fitting our traces^27^ to extract kinetic constants, we consistently observed a rate enhancement for the ligation of the L-over the D-enantiomer, with the enhancements ranging from 5-fold for L-proline to >100-fold for L-lysine and L-leucine (**Figure 2**). In the case of lysine, both the ε- and α-amino groups are potential nucleophiles. We observed similar reactivity for ε-acetyl Lys and Lys, but no reactivity with α-acetyl Lys, indicating that lysyl-RNA ligation occurs exclusively through the α-amine of lysine (**Figure S2**).

**Figure 2.**
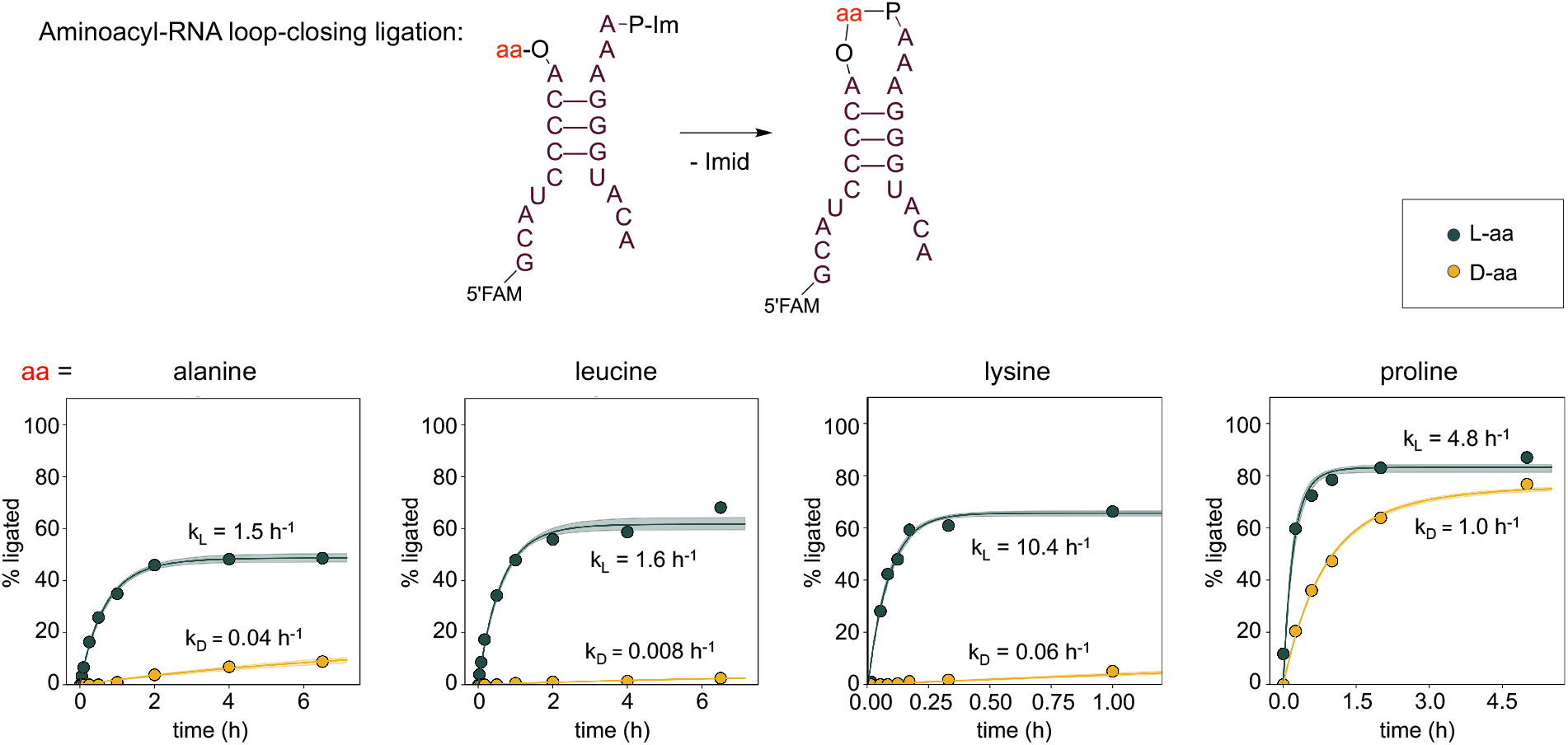
Aminoacyl-RNA loop-closing ligation for RNA architecture 1 with four different amino acids. Top: RNA architecture. Bottom: Time course of loop-closing ligations. The % ligated product was determined as in Figure 1 from gel images. All reactions were conducted in three technical replicates at 0 °C in 5 mM MgCl2, 100 μM Na2EDTA, 100 mM imidazole, pH 8.0 with 5 μM RNA oligonucleotides. Shaded regions envelope the 95% prediction interval as determined from kinetic model.

The loop-closing reaction with RNA architecture 1 is clearly stereoselective for all four amino acids tested. To ask whether the sequence and structure of the RNA plays a role in determining the stereoselectivity of the reaction, we repeated the ligation and hydrolysis experiments with two different RNA architectures (**Figure 3, Figures S4–S5**). In both cases we observed a consistent stereoselective link between D-RNA and L-amino acids (**Figure 3**).

**Figure 3.**
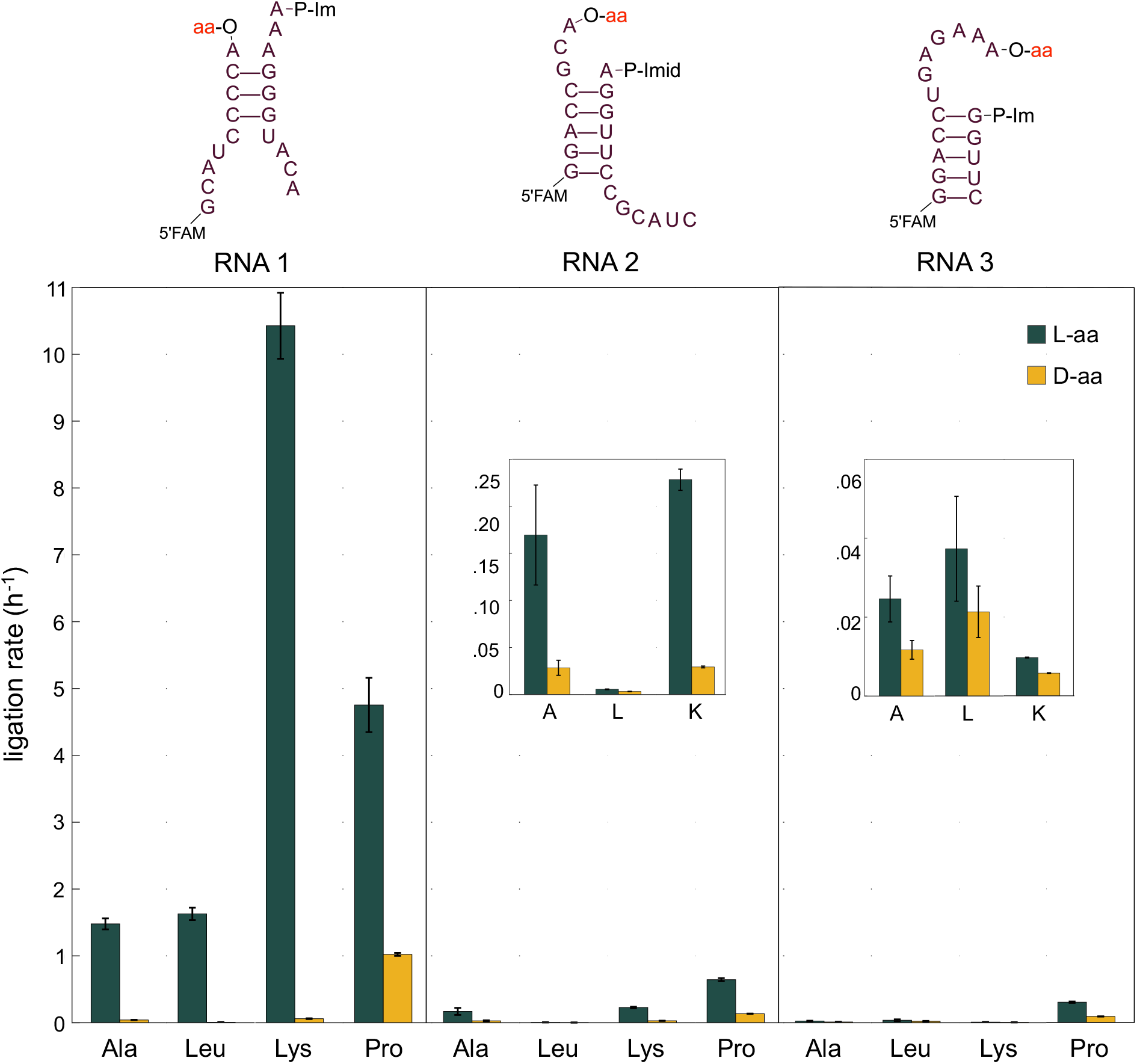
Aminoacyl-RNA ligation rates for three different RNA architectures and four different amino acids. Top: schematics for RNA architectures 1, 2 and 3. Bottom: Observed rates for aminoacyl-RNA loop-closing ligation for the three RNA architectures with the amino acids alanine, leucine, lysine, and proline. Ligation reactions were conducted in triplicate at 0 °C in 5 mM MgCl2, 100 μM Na2EDTA, 100 mM imidazole, pH 8.0, with 5 μM RNA oligonucleotides. Dark green bars: RNA aminoacylated with L-amino acids; yellow bars: RNA aminoacylated with D-amino acids. Ligation rates were derived from kinetic models using ligation yields and experimental rates for aminoacyl hydrolysis (see Methods). All L- and D- pairwise comparisons were statistically significant, with L-aminoacyl loop-closing ligation rates higher than D- in all cases. Error bars and significance were estimated as detailed in Materials and Methods.

However, the magnitude of the stereoselectivity depended strongly on the specific overhang, with RNA 1 exhibiting the greatest stereoselective effect (on the order of 10^1^-10^2^). For RNA 2 and RNA 3, the ligation rate difference between RNA aminocylated with L-vs. D-amino acids ranged from 1.7 fold to ten-fold (**Table 1**). Interestingly, the rates and stereoselectivity of loop-closing ligation varied widely between different combinations of RNA architecture and amino acid, suggesting that specific interactions between the amino acid and the RNA structure influence the outcome of the reaction. Nevertheless, for every combination of overhang and amino acid (12 in total), ligation proceeded more efficiently with L-amino acids than their D-enantiomers (**Figure 3**). Faster rates of loop-closing ligation correlated with higher ligation yields, as expected for a process that is a partition between highly variable ligation and relatively constant hydrolysis (**Figures S3–S6**). In general the hydrolysis reaction exhibited very little stereoselectivity, with a maximum rate difference of 1.6-fold, so that differences in the hydrolysis rates do not explain the observed differences in ligation rates (**Tables S1-S4)**.

**Table 1.** Ligation rate constants for the three RNA architectures with the four amino acids: Ala, Leu, Lys, and Pro. Ligation rates were derived from kinetic models using ligation yields as a function of time and experimental rates for aminoacyl hydrolysis (see Methods). The rightmost column shows the ratio of kL/kD, and all differences between L- and D-ligation rates are statistically significant (p<0.05, non-parametric test).

| | | $k_L$ ($h^{-1}$) | $k_D$ ($h^{-1}$) | $k_L/k_D$ |
| --- | --- | --- | --- | --- |
| RNA 1 | Ala | $1.48 \pm .08$ | $.041 \pm .003$ | 36 |
| | Leu | $1.62 \pm .09$ | $.0084 \pm .0004$ | 194 |
| | Lys | $10.4 \pm .5$ | $.061 \pm .006$ | 172 |
| | Pro | $4.8 \pm .4$ | $1.02 \pm .02$ | 4.7 |
| RNA 2 | Ala | $.17 \pm .06$ | $.028 \pm .008$ | 6.0 |
| | Leu | $.0056 \pm .0003$ | $.0033 \pm .0002$ | 1.7 |
| | Lys | $.23 \pm .01$ | $.029 \pm .001$ | 7.8 |
| | Pro | $.64 \pm .02$ | $.135 \pm .002$ | 4.8 |
| RNA 3 | Ala | $.025 \pm .006$ | $.012 \pm .002$ | 2.1 |
| | Leu | $.037 \pm .013$ | $.021 \pm .007$ | 1.7 |
| | Lys | $.0097 \pm .0001$ | $.0058 \pm .0001$ | 1.7 |
| | Pro | $.30 \pm .01$ | $.091 \pm .001$ | 3.4 |

To determine whether the chirality of the RNA is indeed responsible for the stereoselectivity of the loop-closing ligation reaction, as opposed to any unknown and uncontrolled variable such as the presence of chiral contaminants, we repeated the ligation and hydrolysis reactions using synthetic L-RNA rather than the canonical D-RNA. L-RNA was chemically synthesized using L-nucleotides, and aminoacylated using an L-RNA version of the Flexizyme ribozyme. We selected proline as a test case given that prolyl-RNA ligation proceeded at a high rate with all three architectures of D-RNA. We observed that inversion of the RNA chirality resulted in ligation favoring D-amino acids with all three RNA architectures (**Figure 4)**. While the rate constants were not perfectly mirrored, most likely due to differences in the purity of the synthetic substrates and minor variations in experimental conditions, the overall pattern of the ligation reaction was conserved but reversed between D- and L-RNA. We also conducted an L-RNA reaction using RNA architecture 1 and lysine. In this condition, we observed highly stereoselective ligation with D-lysine over L-lysine (130-fold rate enhancement, **Figure 4, Table 2**).

**Table 2.** Rate constants for loop-closing ligation with aminoacylated L-RNA. Ligation rates were derived from kinetic models using ligation yields as a function of time and experimental rates for aminoacyl hydrolysis (see Methods). The rightmost column shows the ratio of kD/kL, and all differences between L- and D-ligation rates are statistically significant (p<0.05, non-parametric test).

| | | $k_L$ (h<sup>-1</sup>) | $k_D$ (h<sup>-1</sup>) | $k_D/k_L$ |
| --- | --- | --- | --- | --- |
| RNA 1 | Lys | $.067 \pm .001$ | $8.6 \pm .9$ | 130 |
| | Pro | $1.68 \pm .03$ | $11.4 \pm .7$ | 6.8 |
| RNA 2 | Pro | $.52 \pm .02$ | $8.7 \pm .3$ | 17 |
| RNA 3 | Pro | $.114 \pm .005$ | $.24 \pm .01$ | 2.1 |

**Figure 4.**
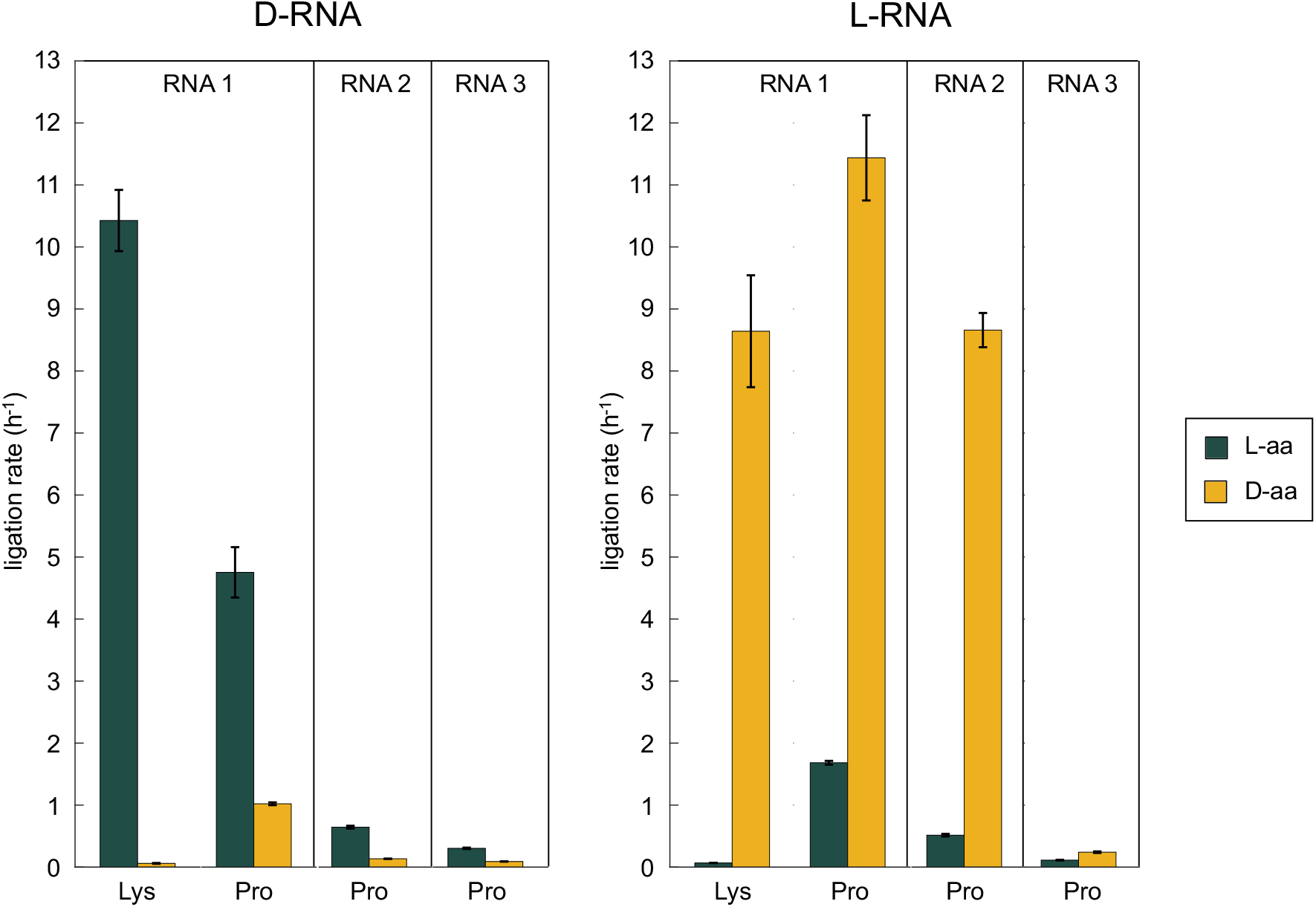
Loop-closing ligation with aminoacylated L-RNA. Rate constants for loop-closing ligation with D-RNA (left) and L-RNA (right). The D- and L-versions of RNA 1 exhibit high stereoselectivity for L- and D-lysine, respectively. Inversion of stereoselectivity is seen for all three D-vs. L-RNAs with proline. Each reaction was conducted in three replicates at 0 °C in 5 mM MgCl2, 100 μM Na2EDTA, 100 mM imidazole, pH 8.0 with 5 μM RNA oligonucleotides. Dark green bars: aminoacylated L-RNA; yellow bars: aminoacylated D-RNA. Error bars and significance (p<0.05) were estimated using the Monte Carlo method as detailed in Materials and Methods.

## Discussion

We have found a consistent bias, across three RNA architectures and with four amino acids, for faster loop-closing ligation when D-RNA is aminoacylated with an L-amino acid as opposed to a D-amino acid. As expected from mirror symmetry, when the L-isomer of RNA is aminoacylated, the ligation reaction proceeds preferentially with D-amino acids. Both the reaction rate for loop-closing ligation and the magnitude of the stereoselectivity are highly dependent on the RNA sequence and structure. Interestingly, higher ligation rates are generally associated with higher stereoselectivity. We suggest that a compact folded aminoacyl-RNA structure that is pre-organized so as to place the amine of the amino acid in proximity to the activated 5′-phosphate would both enhance the reaction rate and lead to greater stereoselectivity. For example, RNA architecture 1, which displays the highest rates of ligation, also exhibits the highest stereoselectivity (**Figure 3**). Architecture 1 differs from the other RNA architectures in that it contains a three nucleotide 5′-overhang and a single nucleotide 3′-overhang, which is aminoacylated. All four overhanging nucleotides are adenosines, which suggests that the strong stacking interactions of the purine nucleobases could lead to a compact folded structure prior to the loop-closing ligation.^28^ Architecture 2 has a single nucleotide 5′-overhang and a putative three nucleotide 3′-overhang; one of the four nucleotides is a pyrimidine while the other three are purines. This architecture led to slower and less stereoselective loop-closing ligation with all four amino acids, for reasons that are unclear. Finally, RNA architecture 3 emerged from a screen for 3′-overhang sequences that led to efficient loop-closing ligation following aminoacylation with glycine. Our structural studies^22^ of the glycine-bridged product structure suggest that its 3′-overhang folds so that it largely envelops the glycine. It is therefore less surprising that this RNA exhibits lower rates of loop-closing ligation, as well as less stereoselectivity than architectures 1 and 2, since aminoacylation with bulkier amino acids may disrupt the folded RNA structure. We hope that future studies will elucidate structural basis for stereoselectivity in the loop-closing reactions that we have studied.

In contrast to ligation, hydrolysis rates of aminoacylated RNA did not vary widely with RNA sequence (**Tables S1-S4**). In all cases, prolyl-RNA hydrolyzed more rapidly than the other aminoacyl-RNA esters, presumably due to the enhanced basicity of the secondary amine. Given that protonation of the amine accelerates hydrolysis of aminoacylated RNA,^29^ we suggest that the relatively small changes in hydrolysis rate between the duplex and single-stranded conditions support the argument that RNA tertiary structure influences the local chemical environment, including the effective pH and steric accessibility, in the vicinity of the aminoacyl amine.

The exclusive use of L-amino acids in biological proteins presents a puzzle. Although a common mechanism that could enrich all 20 amino acids is possible, it has eluded discovery.

The concept of chiral propagation is an elegant solution to the problem of the homochirality of proteinogenic amino acids in biology, but the molecular details behind this process remain obscure. Ozturk et al,^30^ following the argument by Tamura and Schimmel,^31^ have posited that the homochirality of RNA was established first, followed by diastereoselective aminoacylation allowing for synthesis of homochiral peptides from racemic amino acids, and subsequently, chiral transfer processes dictated the handedness of biological metabolites. Alternatively, one might imagine that primordial metabolic reactions catalyzed by ribozymes would begin to fix the chirality of metabolic intermediates, including amino acids, due to the intrinsic stereoselectivity of macromolecular catalysis. However, this leaves unanswered the question of whether there is some underlying reason that D-RNA would lead ultimately to the coded synthesis of peptides and proteins composed exclusively of L-(and not D-) amino acids. Considering the earlier reports from Tamura and Schimmel,^17^ Roberts et al.,^19^ and Radakovic et al.,^21^ it appears that L-amino acid selectivity arises readily in reactions involving aminoacylated RNA. Our results with loop-closing ligation mediated by aminoacylated RNA confirm and extend this connection between D-RNA and L-amino acids. We have previously demonstrated that loop-closing ligation with aminoacylated RNAs can lead to the assembly of functional ribozymes.^25^ The resulting ribozymes are composed of RNA segments bridged by aminoacylester phosphoramidates. We have proposed^25^ that this process, operating in primordial protocells, may have facilitated the assembly of ribozymes from smaller RNA fragments that could have been replicated by nonenzymatic chemistry. Due to the stereoselectivity of loop-closing ligation with aminoacylated RNA, these chimeric ribozymes would largely contain bridging L-amino acids. We hypothesize that in such chimeric ribozymes, the amino acid side chain may have assisted in catalysis, leading to a selection pressure for the emergence of stereoselective aminoacylating ribozymes capable of putting specific amino acids on specific RNAs. The emergence of RNA aminoacylating ribozymes with specificity for L-amino acids would then have provided the evolutionary foundation for the later synthesis of homochiral proteins from L-amino acids. This scheme, while hypothetical, presents an experimentally tractable system for investigating the origin of amino acid homochirality.

## TOC Figure

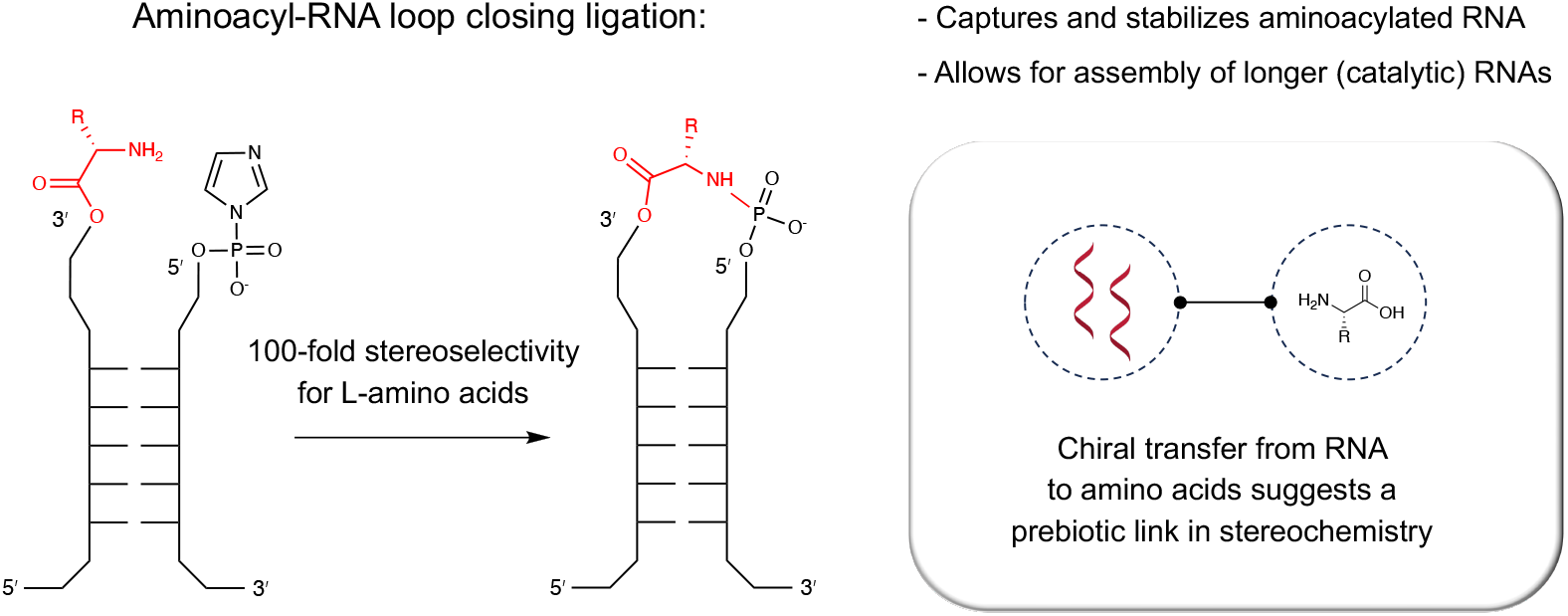

Summary:

In this work, we test for potential stereoselectivity in aminoacyl-RNA loop closing ligation. We find that the reaction preferentially captures L-amino acids, with implications towards the origin of amino acid homochirality.

