## Supplementary Information for "Stereoselectivity of Aminoacyl-RNA Loop-closing Ligation" (from another version of this preprint: version 1, 2024-11-27)

Table of Contents

1. Materials and Method ........................................................ S2
2. Supplementary Tables ........................................................ S8
3. Supplementary Figures ........................................................ S11

#### Materials and Methods

Oligonucleotide Synthesis

Stem-loop oligonucleotides were prepared on a K&A H-8 oligo synthesizer with RNA amidites purchased from Chemgenes (part no. ANP-5671, ANP-6676, ANP-5673, ANP-5674, CLP-1454). Amidites were dissolved at 67 mM in acetonitrile, and synthesis (including removal of DMT) was carried out on a 1 μmol scale controlled-pore-glass (CPG) column. Following synthesis, oligonucleotides were cleaved from the column in 1.2 mL of 1:1(v/v) $NH_4OH$:methylamine (AMA); the solution was passed 20 times bidirectionally through the column, and the submerged column was incubated for 15 minutes at room temperature. The solution and an additional 1 mL wash were collected and incubated at 65 °C for 15 minutes. The samples were then evaporated by SpeedVac followed by freeze drying. The dried samples were redissolved in 100 μL DMSO and combined with 125 μL triethylamine trihydrofluoride (TEA.3HF) to deprotect the 2′-hydroxyl groups on the RNA backbone. The deprotection reaction was heated at 65 °C for 2.5 hours, then precipitated by addition of 22.5 μL 5 M ammonium acetate and 1 mL isopropanol, incubation on dry ice for 15 minutes and spun down at 16,000xg for 15 min, the supernatant was discarded, and the same procedure was repeated with 1 mL 80 %(v/v) ethanol. Ribozymes were prepared using the K&A synthesizer or by *in vitro* transcription of a DNA template (New England BioLabs HiScribe T7 Quick High Yield RNA Synthesis kit, DNA purchased from IDT). L RNA oligonucleotides were prepared using the K&A synthesizer with L-amidites purchased from Chemgenes (part no. ANP-4841, ANP-4846, ANP-4843, and ANP-4845). All oligonucleotides, following synthesis or *in vitro* transcription, were purified on a 20 % (v/v) Urea-PAGE gel, desalted with Sep-Pack C18 column and verified mass with LC-MS.

Oligonucleotide Aminoacylation

Aminoacylated oligonucleotides were prepared using modified versions of the Flexizyme, an aminoacyl-transfer ribozyme. We used the dFx Flexizyme[26] to aminoacylate oligo 3A, and two derivative Flexizymes which were modified at the 3′ terminus to allow base pairing with oligos 1A and 2A (all sequences may be found in Table S5). 3,5-dinitrobenzyl esters of amino acids (aa-DBEs), the substrate for Flexizyme aminoacylation, were prepared as described below. Aminoacylation reactions were carried out at 0 °C in a total solution volume of 500 μL. Each reaction contained 100 mM imidazole pH 8, 5 mM $MgCl_2$, 30 μM oligonucleotide, 30 μM Flexizyme, and 5 mM aa-DBE (20 % DMSO v/v). Reactions were incubated at 0 °C for 1-24 hours depending on the amino acid: proline 1 hr, alanine 3-6 hr, leucine 6-12 hr, lysine 12-24 hr. Upon completion, reactions were concentrated to 100 μL by centrifugal filtration (Amicon 0.5 mL 3K MWCO, 10 minutes at 21,300 x g), then precipitated by addition of 40 μL of 3 M sodium acetate pH 5 and excess alcohol (ethanol or isopropanol). Pellets were washed twice with 80 % ethanol and dried under vacuum. These samples were redissolved in 100 μL 99 % v/v formamide with 5 mM $Na_2$EDTA and purified on an acidic 20 % v/v Urea-PAGE gel (80 mL gel volume, poured and run with 100 mM sodium acetate buffer, pH 5.0) to separate aminoacylated oligonucleotides from starting material and Flexizyme RNA. Aminoacylated bands were cut, crushed, and extracted in a 1.5 mL pH 5.0 buffer with 50 mM NaOAc and 5 mM EDTA buffer for three hours while rotating at 4°C. After extraction, the solution was separated from gel particles by filtering through 5 µm filter and RNA was precipitated by addition of sodium acetate and ethanol, washed twice with 80% ethanol, and redissolved in 100 µL water. The product was flash frozen on dry ice and stored at -80°C.

3,5-Dinitrobenzyl Ester (DBE) Amino Acid Synthesis

DBE amino acids were prepared by combining 0.6 mmol of the Boc-protected amino acid with 1 mmol triethylamine, and 0.5 mmol 3,5-dinitrobenzyl chloride in 150 µL of N,N-dimethylformamide. The reaction was stirred at room temperature overnight, then transferred to a separatory funnel with 10 mL of diethyl ether. The product was washed three times with 0.5 M HCl, then washed three times with 4% $NaHCO_3$, then washed once with saturated NaCl. The remaining organic phase was dried with solid $MgSO_4$, vacuum filtered, and evaporated using a rotary evaporator. The solid product (ranging from off-white powder to yellow-orange oil, depending on the amino acid) was transferred to a glass tube and deprotected by addition of 2 mL trifluoroacetic acid (TFA) and incubation at room temperature for 10 minutes. TFA was removed under a nitrogen stream, and the remaining product was washed three times with diethyl ether, with rotary evaporation after each wash. The final product was triturated with diethyl ether, dried under nitrogen, then dissolved in DMSO. The correct product was confirmed by NMR (400 MHz).

Oligonucleotide Activation with DPDS and Imidazole

To activate the 5′-phosphate of capture-strand oligonucleotides with imidazole: the dry RNA was dissolved in 100 µL DMSO, followed by addition of 2.7 mg imidazole and 5.3 mg triphenyl phosphine (TPP). After complete dissolution, 7 µL triethylamine (TEA) was added, followed by 4.4 mg of 2,2'-dipyridyldisulfide (DPDS) as the coupling agent. The reaction was incubated with rotation at room temperature for three hours. After three hours, the same amounts of imidazole, TPP, TEA and DPDS were added, and the reaction was incubated overnight. The next day, the RNA was precipitated by addition of 100 µL saturated sodium perchlorate in acetone and 1 mL acetone, then washed with 1:1 v/v acetone:diethylether. The level of activation of each oligonucleotide was measured by high-performance liquid chromatography (HPLC) with an Agilent ZORBAX analytical column (Eclipse Plus C18, 250 × 4.6 mm, 5-µm particle size, 959990-902) at a flow rate of 1 mL/min. The separation gradients were aqueous 20 mM triethylammonium bicarbonate, pH 8.0 (A), and acetonitrile (B), from 6 to 14% B over 24 min for RNAs 1 and 2 and from 6 to 11% B over 24 minutes for RNA 3. The measured values for percent imidazole-activation were: 83% for RNA 1, 65% for RNA 2, 84% for RNA 3.

Oligonucleotide activation:

| Reagent | Amount |
|---|---|
| 5’ Phosphorylated RNA | Up to 0.001 mmol |
| DMSO | 100 μL |
| Imidazole | 0.040 mmol |
| TPP | 0.020 mmol |
| TEA | 0.050 mmol |
| DPDS | 0.020 mmol |

<u>Ligation and Hydrolysis Reactions</u>

Reagents were added to PCR tubes to the final concentrations tabulated below. Reactions were set up in three parallel replicates, and each reaction volume totaled to 10 μL. Water, MgCl2, imidazole pH 8, and Na2EDTA were added first, and the mixture was equilibrated on ice. Then, the imidazole-activated oligonucleotide (in the case of ligation), 5′-phosphorylated oligonucleotide (in the case of duplex hydrolysis), or water (in the case of single-stranded hydrolysis) was added, followed by the addition of the 3′-aminoacylated oligo. An initial sample of each reaction was taken in quenching buffer (see below), and a time course was initiated. Subsequent time-course samples were diluted with quenching buffer as follows. Ligation reactions were quenched in 92% formamide with 20 mM Na2EDTA and bromophenol blue dye. Hydrolysis reactions were quenched in 70% formamide with 10 mM Na2EDTA, 100 mM sodium acetate pH 5.5, 150 mM HCl, and bromophenol blue dye. The quenched reactions were run on Urea-PAGE gels (40 mL gel volume). Ligation gels were poured and run with 1X TBE buffer, whereas hydrolysis gels were poured and run with 100 mM sodium acetate buffer, pH 5.0. Gels were visualized with the Amersham Typhoon Biomolecular Imager, scanning with the Cy2 wavelength settings. Band intensity was quantified using ImageQuant software and plotted in Excel.

Loop-Closing Ligation reaction mixtures:

| Reagent | Concentration |
|---|---|
| Imidazole pH 8 | 100 mM |
| $MgCl_2$ | 5 mM |
| $Na_2EDTA$ | 100μM |
| 3′ Aminoacylated RNA | 5 μM |
| 5′ Activated RNA | 5 μM |

Duplex hydrolysis reaction mixtures:

| Reagent | Concentration |
|---|---|
| Imidazole pH 8 | 100 mM |
| $MgCl_2$ | 5 mM |
| $Na_2EDTA$ | 100μM |
| 3′ Aminoacylated RNA | 5 μM |
| 5′ phosphorylated RNA | 5 μM |

Single-stranded hydrolysis reaction mixtures:

| Reagent | Concentration |
|---|---|
| Imidazole pH 8 | 100 mM |
| $MgCl_2$ | 5 mM |
| $Na_2EDTA$ | 100μM |
| 3′ Aminoacylated RNA | 5 μM |

Kinetic Analysis

All kinetic rates have been estimated by non-linear least squared curve fitting of time traces using the Levenberg-Marquardt algorithm.[27] For single-stranded hydrolysis data (adjusted R2 > 0.830, median 0.992) and duplex hydrolysis data (adjusted R2 > 0.944, median 0.995), a simple exponential decay model has been employed, with the hydrolysis traces fit as $Ae^{-kt}$, with $k$ being the hydrolysis rate and $A$ an amplitude parameter used to account for small deviations in the estimates of the starting material, typically larger than 0.95.

Ligation reactions have been modeled as parallel irreversible reactions to capture the competition between ligation and hydrolysis (adjusted $R^2 > 0.965$, median 0.991). The following system of differential equations was numerically integrated and used to model the reaction:

$$\begin{cases} d[O1_{AA}]/dt = -[O1_{AA}]k_{h,AA} - [O1_{AA}\!:O2_{IM}]k_{lig} \\ d[O1_{OH}]/dt = [O1_{AA}]k_{h,AA} \\ d[O2_{IM}]/dt = -[O2_{IM}]k_{h,IM} - [O1_{AA}\!:O2_{IM}]k_{lig} \\ d[O2_{OH}]/dt = [O2_{IM}]k_{h,IM} \\ d[O1O2]/dt = [O1_{AA}\!:O2_{IM}]k_{lig} \end{cases}$$

With O1 being the aminoacylated oligonucleotide, O2 its imidazole-activated partner and O1:O2 their non-covalent complex that can be formed with any combination of $O1_{OH}$, $O1_{AA}$, $O2_{OH}$, $O2_{IM}$. Importantly, only the duplex bearing both aminoacyl- and imidazole- moieties can undergo ligation and produce a covalently linked species O1O2.

The aminoacyl-RNA hydrolysis ($k_{h,\,AA}$) used in the model was as determined from our experiments measuring hydrolysis in a duplex, and the 5′-phosphorimidazolide hydrolysis ($k_{h,\,IM}$) used in the model was as determined from our recent work.

The model was numerically integrated using Euler's method and dt ≈ 1 min, and the reshuffling of oligonucleotides in the O1:O2 complexes ($O1_{OH}$:$O2_{OH}$, $O1_{AA}$:$O2_{OH}$, $O1_{OH}$:$O2_{IM}$, $O1_{AA}$:$O2_{IM}$) was updated at every time step under the assumption of a fast equilibration rate, with the reactive complex concentration [$O1_{AA}$:$O2_{IM}$] computed as follows:

$$[O1_{AA}\!:O2_{IM}] = \frac{[O1_{AA}]}{[O1_{AA}] + [O1_{OH}]} * \frac{[O2_{IM}]}{[O2_{IM}] + [O2_{OH}]} * ([O1_{AA}] + [O1_{OH}])$$

The ligation rate ($k_{lig}$) and the initial purity of the imidazole-activated O2 ($O2_{IM}/O2_{tot}$) were left as free parameters to be fitted on the experimental data.

In some cases, the percentage of initial imidazole-activation estimated from the kinetic model was noticeably low. We analyzed the percent imidazole-activation by HPLC after the conclusion of kinetic experiments, and observed % activation of 85%, 65%, and 85% for RNAs 1, 2, and 3, respectively. While most of the percent-activation values indicated by the kinetic model did fall within the 60-100% range, a handful did not (**Figure S4**). Other influences, such as degradation of the RNA itself, might also be contributing to this underestimation. However, any such effects should equally impact the L- and D-aminoacylated reactions, as the capture strand RNA was identical between them in all experiments.

Statistical Analysis

Due to heteroscedasticity of our data (lower variance as we approach zero and one), the standard errors on the coefficients as provided from the nonlinear curve fitting algorithm cannot be used reliably. To estimate errors associated with our rate constants, we used the Monte Carlo method to generate synthetic time traces, with every data point sampled from a normal distribution having mean and standard deviation determined experimentally from three independent replicates. Every set of generated datapoints was independently fitted and the average and standard deviations of the resulting coefficients are reported in this work.

Since the specific amino acid and RNA sequence identity heavily affect the reactions, to determine whether amino acid chirality has an effect on the reaction rates, we had to compare them on a case-by-case basis. Briefly, for each iteration of the Monte Carlo routine to generate a new sample and determine the associated rate constants, a comparison with a paired sample of opposite chirality was performed and the differences between the rates ($\Delta k$) were stored. To reject or accept the null-hypothesis of equivalence of the rates in the two conditions ($H_0$: $\Delta k = 0$), we performed a non-parametric test to check whether the null value was included in the 95% confidence interval of our distributions.

#### Supplementary Tables

**Table S1**. Hydrolysis rates for aminoacyl-RNA annealed to the capture strand ("duplex hydrolysis"), for three RNAs with the four amino acids alanine, leucine, lysine, and proline. Reactions were conducted in triplicate at 0 °C in 5 mM $MgCl_2$, 100 µM $Na_2EDTA$, 100 mM imidazole, pH 8.0 with 5 µM oligonucleotides. The rightmost column shows the ratio of $k_L/k_D$, and asterisks* indicate a statistically significant rate difference ($p<0.05$, non-parametric test).

| | | $k_L$ ($h^{-1}$) | $k_D$ ($h^{-1}$) | $k_L/k_D$ |
|---|---|---|---|---|
| RNA 1 | Ala | .097 ± .001 | .083 ± .001 | 1.2* |
| | Leu | .070 ± .001 | .100 ± .003 | 0.7* |
| | Lys | .055 ± .001 | .0514 ± .0003 | 1.1* |
| | Pro | .22 ± .02 | .21 ± .02 | 1.1 |
| RNA 2 | Ala | .18 ± .01 | .17 ± .02 | 1.0 |
| | Leu | .096 ± .003 | .107 ± .002 | 0.9* |
| | Lys | .108 ± .002 | .10 ± .01 | 1.1 |
| | Pro | .55 ± .02 | .65 ± .01 | 0.8* |
| RNA 3 | Ala | .043 ± .008 | .062 ± .006 | 0.7 |
| | Leu | .050 ± .004 | .060 ± .003 | 0.8* |
| | Lys | .021 ± .002 | .028 ± .003 | 0.7* |
| | Pro | .16 ± .01 | .29 ± .06 | 0.6* |

**Table S2**. Hydrolysis rates for aminoacyl-RNA in the absence of the capture strand ("single-stranded hydrolysis"), for three RNA constructs with the four amino acids alanine, leucine, lysine, and proline. Reactions were conducted in triplicate at 0 °C in 5 mM $MgCl_2$, 100 µM $Na_2EDTA$, 100 mM imidazole, pH 8.0 with 5 µM oligonucleotide. The rightmost column shows the ratio of $k_L/k_D$, and asterisks* indicate a statistically significant rate difference ($p<0.05$, non-parametric test).

| | | $k_L$ ($h^{-1}$) | $k_D$ ($h^{-1}$) | $k_L/k_D$ |
|---|---|---|---|---|
| RNA 1 | Ala | .136 ± .001 | .130 ± .002 | 1.0 |
| | Leu | .075 ± .002 | .114 ± .003 | 0.7* |
| | Lys | .065 ± .001 | .064 ± .001 | 1.0 |
| | Pro | .25 ± .01 | .24 ± .01 | 1.1 |
| RNA 2 | Ala | .116 ± .002 | .118 ± .001 | 1.0* |
| | Leu | 7.7 ± 0.4 | .11 ± .01 | 0.7* |
| | Lys | .054 ± .001 | .049 ± .002 | 1.1* |
| | Pro | .33 ± .01 | .31 ± .01 | 1.1 |
| RNA 3 | Ala | .044 ± .10 | .058 ± .003 | 0.8 |
| | Leu | .036 ± .003 | .044 ± .001 | 0.8* |
| | Lys | .0301 ± .0002 | .0343 ± .0004 | 0.9* |
| | Pro | .12 ± .02 | .16 ± .02 | 0.8 |

**Table S3**. Hydrolysis rates for aminoacyl-RNA annealed to the capture strand ("duplex hydrolysis"), for L-RNA. Reactions were conducted in triplicates at 0 °C in 5 mM $MgCl_2$, 100 μM $Na_2EDTA$, 100 mM imidazole, pH 8.0 with 5 μM oligonucleotide. The rightmost column shows the ratio of $k_D/k_L$, and asterisks* indicate a statistically significant rate difference ($p<0.05$, non-parametric test).

| | | $k_L$ ($h^{-1}$) | $k_D$ ($h^{-1}$) | $k_D/k_L$ |
|---|---|---|---|---|
| RNA 1 | Lys | .060 ± .001 | .067 ± .003 | 1.1* |
| | Pro | .16 ± .01 | .18 ± .02 | 1.1 |
| RNA 2 | Pro | .88 ± .11 | .78 ± .01 | 0.9 |
| RNA 3 | Pro | .28 ± .02 | .13 ± .01 | 0.5* |

**Table S4**. Hydrolysis rates for aminoacyl-RNA in the absence of the capture strand ("single-stranded hydrolysis"), for L-RNA. Reactions were conducted in triplicates at 0 °C in 5 mM $MgCl_2$, 100 μM $Na_2EDTA$, 100 mM imidazole, pH 8.0 with 5 μM oligonucleotide. The rightmost column shows the ratio of $k_D/k_L$, and asterisks* indicate a statistically significant rate difference ($p<0.05$, non-parametric test).

| | | $k_L$ ($h^{-1}$) | $k_D$ ($h^{-1}$) | $k_D/k_L$ |
|---|---|---|---|---|
| L-RNA 1 | Lys | .076 ± .003 | .074 ± .003 | 1.0 |
| | Pro | .18 ± .02 | .22 ± .03 | 1.2 |
| L-RNA 2 | Pro | .34 ± .01 | .34 ± .01 | 1.0 |
| L-RNA 3 | Pro | .15 ± .01 | .14 ± .01 | 0.9 |

**Table S5: Oligonucleotides used in this study**

| Name | Sequence | Use |
|---|---|---|
| 1A | 5′FAM-GCAUCCCCA | Aminoacylated oligonucleotide 1 |
| 1B* | 5′Imid-AAAGGGUACA | Ligation with 1A |
| 1B | 5′ -AAAGGGUACA | Duplex hydrolysis with 1A |
| 1F | 5′-GGAUCGAAAGAUUUCCGCAUCCCCGAAAGGGUACAUGGCGUUAGGU | Aminoacylation of 1A |
| 2A | 5′FAM-GGACCGCA | Aminoacylated oligonucleotide 2 |
| 2B* | 5′Imid-AGGUUCCGCAUC | Ligation with 2A |
| 2B | 5′-AGGUUCCGCAUC | Duplex hydrolysis with 2A |
| 2F | 5′-GGACCGCAAGGUUCCGCAUCCUCACGCAAGUGGGUACAUGGCGUUAGCG | Aminoacylation of 2A |
| 3A | 5′FAM-GGACCUGAGAAA | Aminoacylated oligonucleotide 3 |
| 3B* | 5′Imid-GGUUC | Ligation with 3A |
| 3B | 5′-GGUUC | Duplex hydrolysis with 3A |
| 3F | 5′-GGACCGCAAGGUUCCGCAUCCUCAUCGCAAGGUGGGUACAUGGCGUUAUUCUC | Aminoacylation of 3A |

#### Supplementary Figures

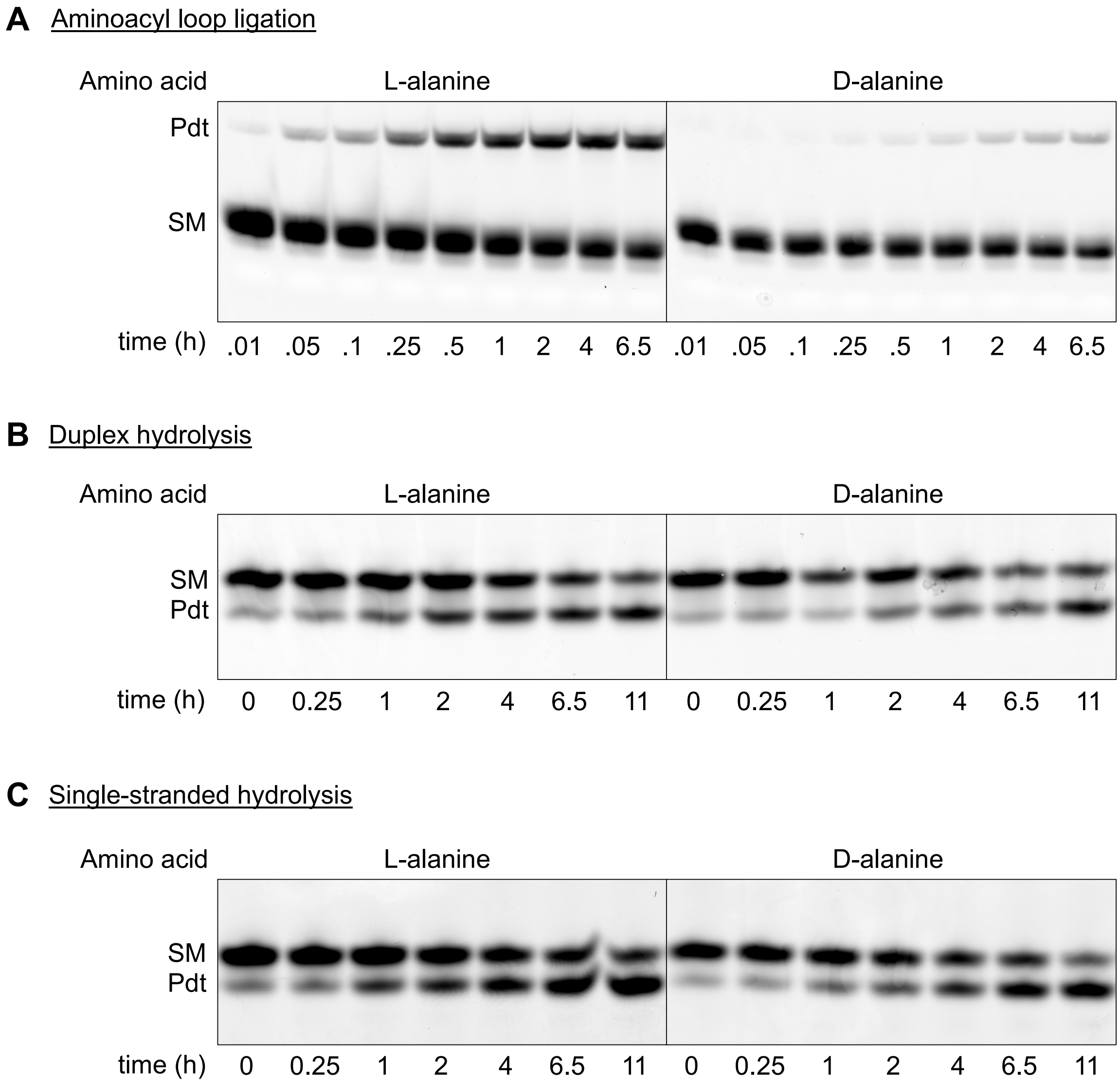


**Figure S1**. **Scans of urea-polyacrylamide gel electrophoresis for reactions of RNA 1 with L- and D-alanine. A.** Aminoacyl-RNA loop closing ligation, where the top band is the aminoacyl-ligated product and the bottom band is the non-ligated starting material. Product band intensity is greater with L-proline than with D-proline. **B.** Hydrolysis of aminoacyl-RNA in a base-paired complex, where the top band is aminoacylated RNA and bottom band is the hydrolysis product. **C.** Hydrolysis of the aminoacyl-RNA in a single-stranded condition, where the top band is aminoacylated RNA and bottom band is the hydrolysis product. Ligation gels were run at pH 8.3, and hydrolysis gels were run at pH 5. All scans were generated with the Amersham Typhoon Biomolecular Imager as described in Methods.

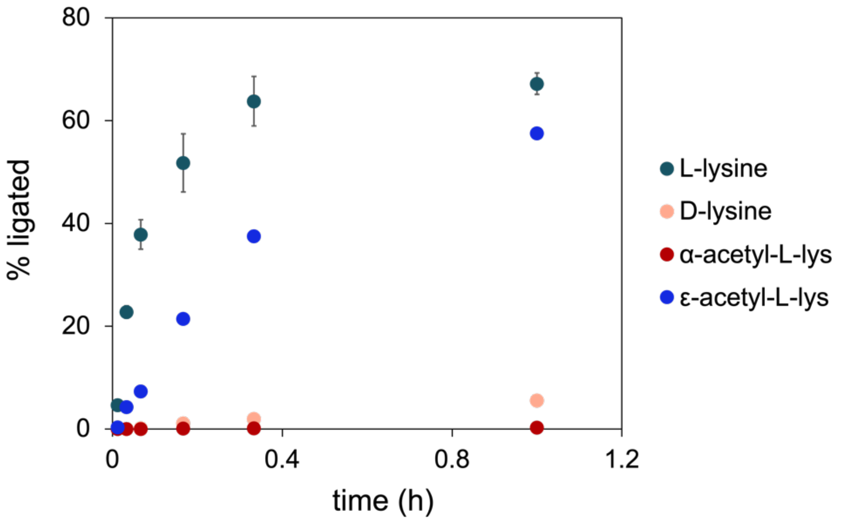


**Figure S2**. **Stereo- and regio-selectivity of loop-closing ligation with lysine.** Time course of loop-closing ligation with RNA 1 and L-lysine, D-lysine, α-acetyl lysine and ε-acetyl lysine. α-acetylated lysine results in near-zero product, whereas ε acetylated lysine ligates efficiently, consistent with the reaction proceeding through attack of the α-amino group on the activated 5′-phosphate.

RNA 1, alanine:

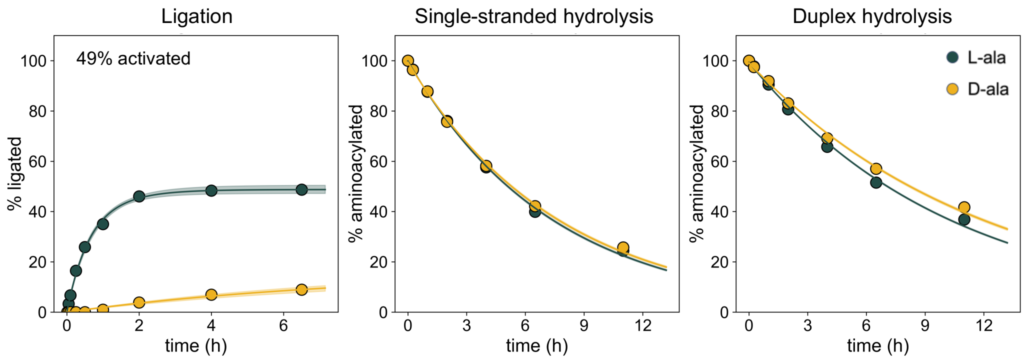


RNA 1, leucine:

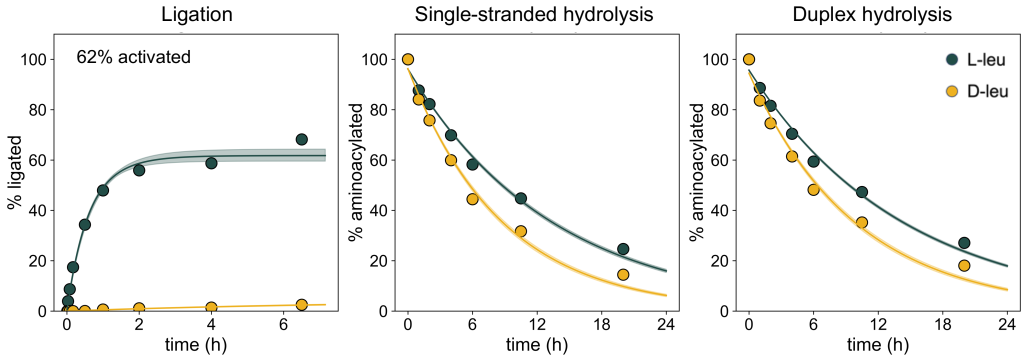


RNA 1, lysine:

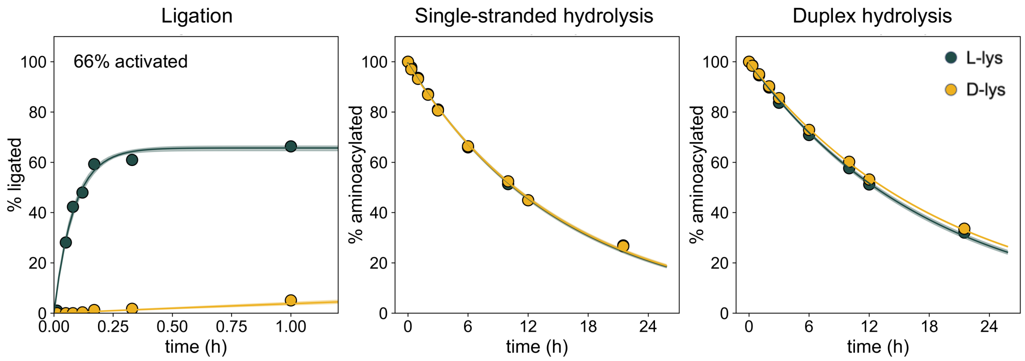


RNA 1, proline:

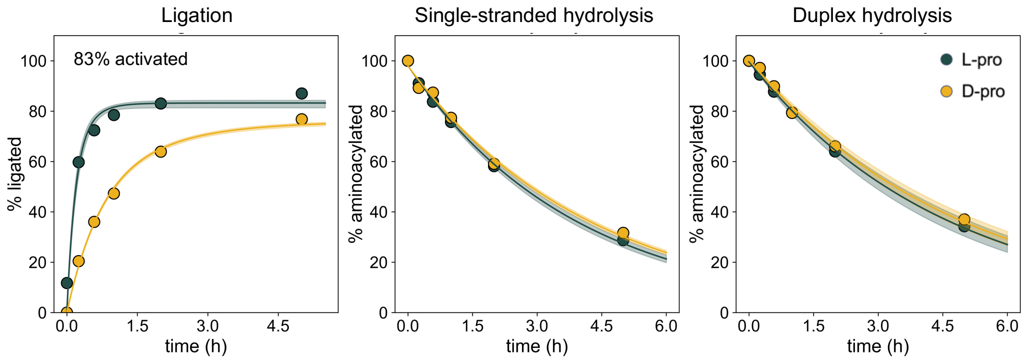

**Figure S3**. **Time courses of loop-closing ligation and hydrolysis for RNA 1.** Values are normalized to the fraction of oligonucleotide 1A aminoacylated at time zero. The percent of capture strand that is imidazole-activated, as estimated from our kinetic models, is stated on the upper right of the plots for ligation. Points show collected data points (average of triplicates) and overlaid shaded areas envelope the empirical 95% prediction interval. Data shown in yellow shows results with L-aminoacylated RNA, and data shown in green D-aminoacylated RNA.

RNA 2, alanine:

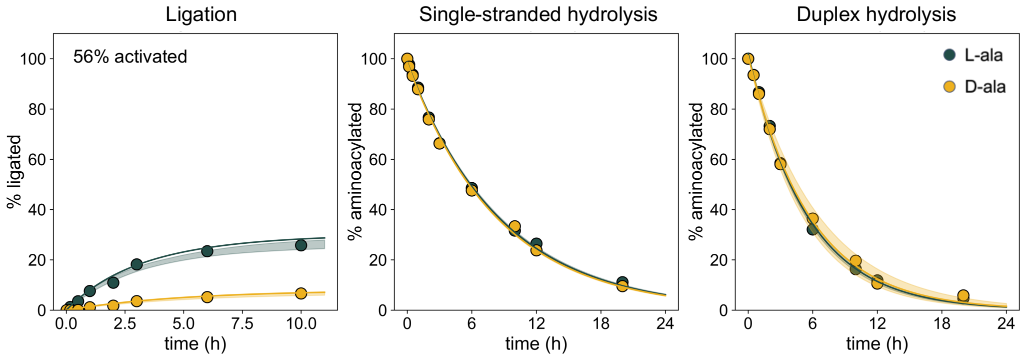


RNA 2, leucine:

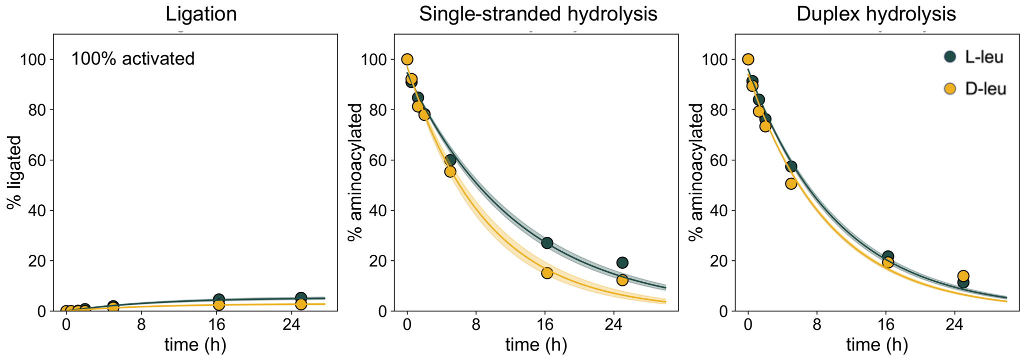


RNA 2, lysine:

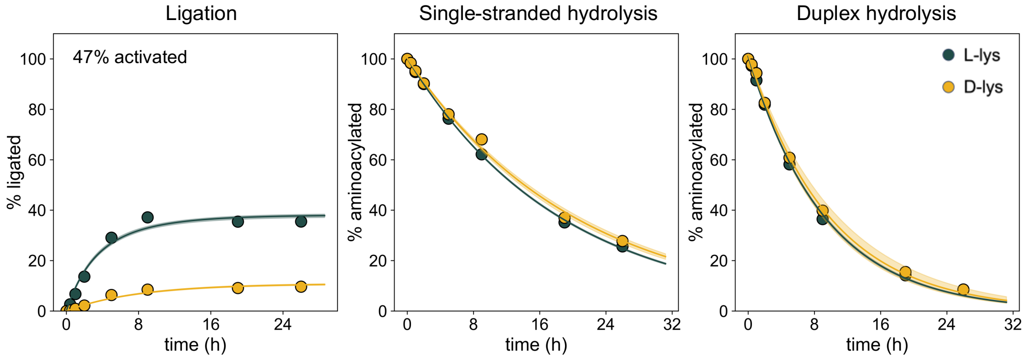


RNA 2, proline:

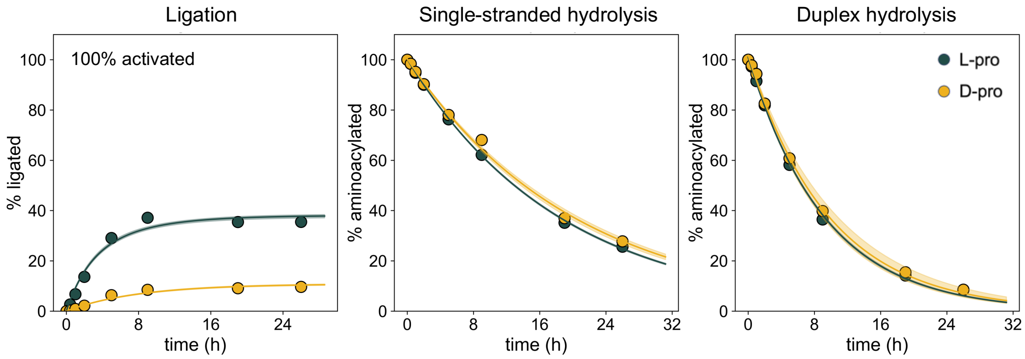

**Figure S4**. **Time courses of loop-closing ligation and hydrolysis for RNA 2.** Values are normalized to the portion of oligonucleotide 2A aminoacylated at time zero. The imidazole-activated percent of capture strand predicted by kinetic models is stated on the plot for ligation. Points show collected data points (average of triplicates) and shaded areas envelope the empirical 95% prediction interval. Data shown in yellow shows results with L-aminoacylated RNA, and data shown in green D-aminoacylated RNA.

RNA 3, alanine:

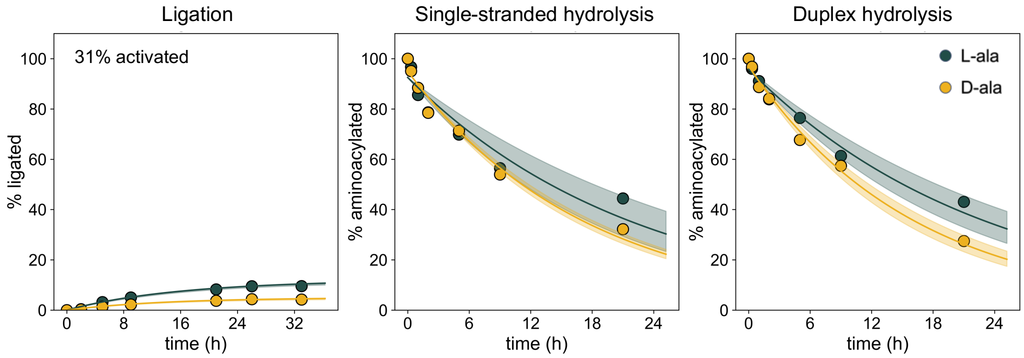


RNA 3, leucine:

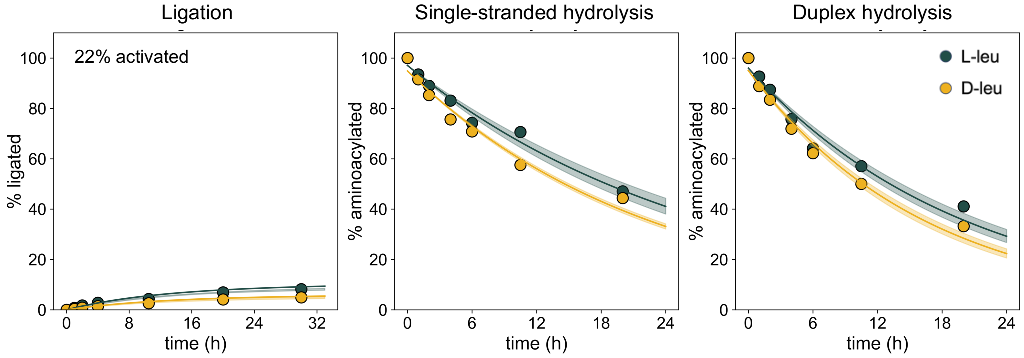


RNA 3, lysine:

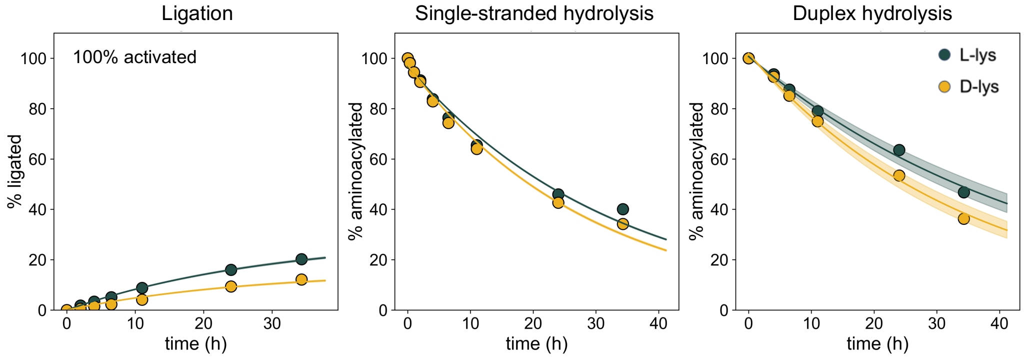


RNA 3, proline:

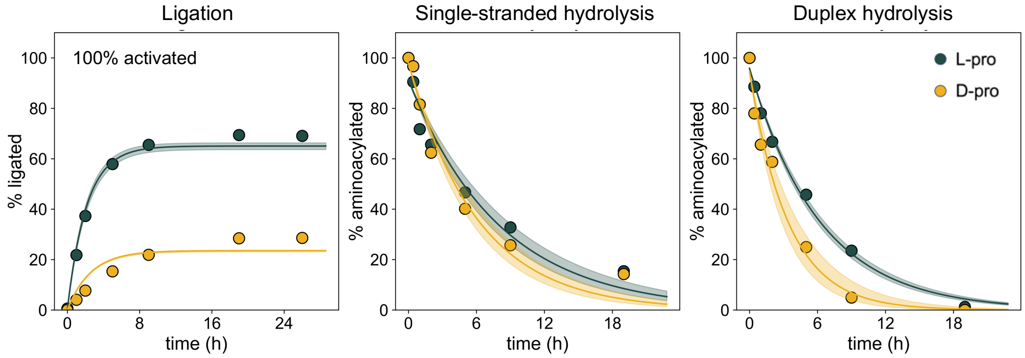

**Figure S5. Time courses of loop-closing ligation and hydrolysis for RNA 3.** Values are normalized to the portion of oligonucleotide 3A aminoacylated at time zero. The imidazole-activated percent of capture strand predicted by kinetic models is stated on the plot for ligation. Points show collected data points (average of triplicates) and shaded areas envelope the empirical 95% prediction interval. Data shown in yellow shows results with L-aminoacylated RNA, and data shown in green D-aminoacylated RNA.

### L-RNA 1, lysine:

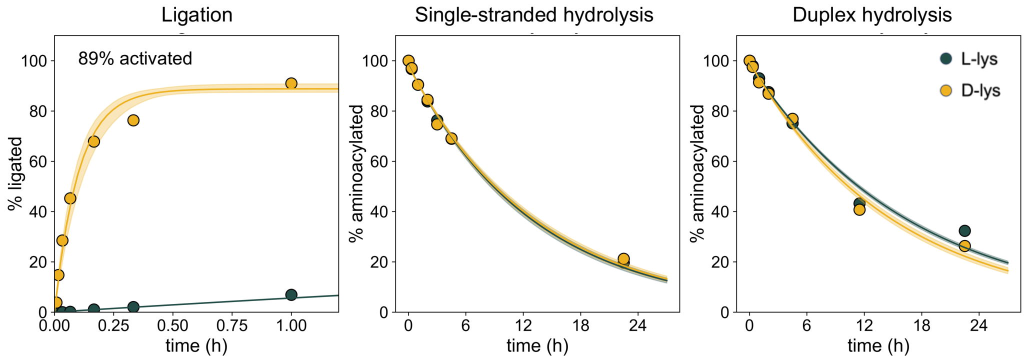


### L-RNA 1, proline:

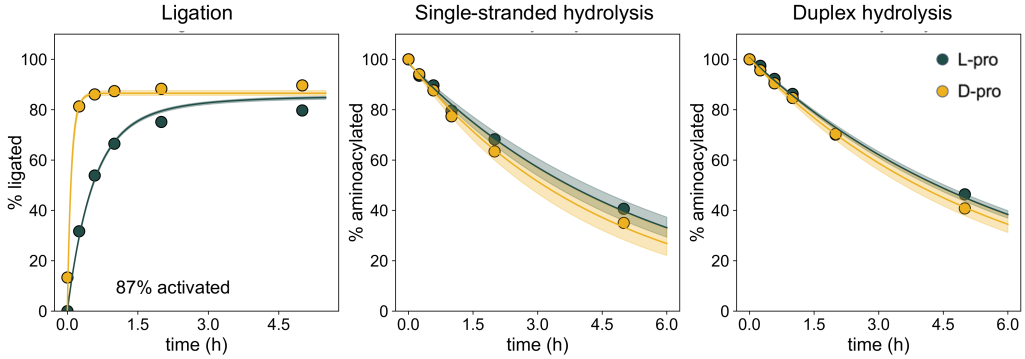


### L-RNA 2, proline:

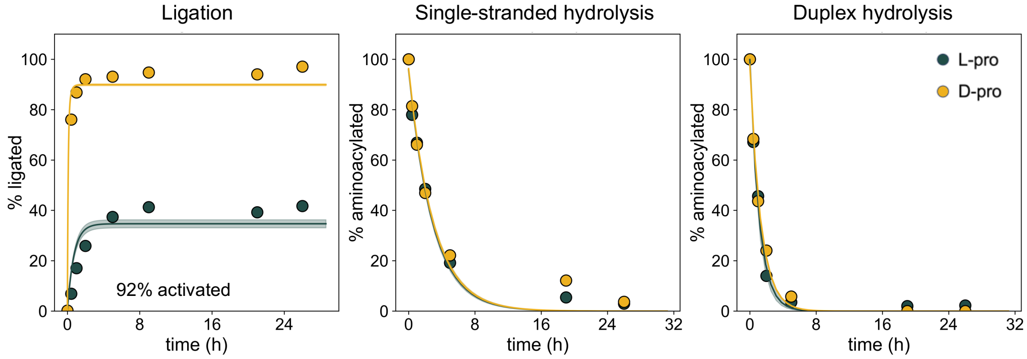


### L-RNA 3, proline:

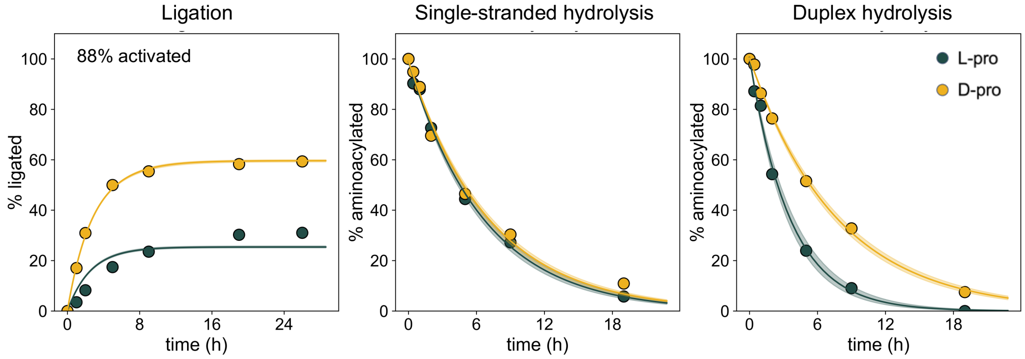

**Figure S6**. **Time courses of loop-closing ligation and hydrolysis for L-RNAs 1, 2 and 3 with proline.** Values are normalized to the portion of oligonucleotides 1A, 2A, and 3A aminoacylated at time zero. The imidazole-activated percent of capture strand predicted by kinetic models is stated on the plot for ligation. Points show collected data points (average of triplicates) and shaded areas envelope the empirical 95% prediction interval. Data shown in yellow shows results with L-aminoacylated RNA, and data shown in green D-aminoacylated RNA.
